# Sequencing and annotation of the endangered wild buffalo (*Bubalus arnee*) mitogenome for taxonomic and hybridization assessment

**DOI:** 10.1101/2020.05.19.103952

**Authors:** Ankit Shankar Pacha, Parag Nigam, Bivash Pandav, Samrat Mondol

## Abstract

The wild water buffalo (Bubalus arnee) is one of the most endangered and least studied large bovid in the Indian subcontinent. India retains 90% of the global population as two fragmented populations in Assam and Chhattisgarh, both threatened by habitat loss and degradation, hunting, disease from livestock, and hybridization with the domestic buffalos. For the first time, we sequenced the 16,357 bp long mitogenome of pure wild water buffalo from both populations. The annotated genes included 13 protein-coding genes, 22 tRNA, two ribosomal genes, and a non-coding control region. Comparative mitogenome analyses showed both populations are genetically similar but significantly different from domestic buffalo. We also identified structural differences in seven tRNA secondary structures between both species. Both wild and domestic water buffalo formed sister clades which were paraphyletic to genus Bos. This study provides baseline information on wild buffalo mitogenome for further research on phylogeny, phylogeography and hybrid assessment and help conserving this endangered species.

## 1. Introduction

The wild water buffalo (*Bubalus arnee*) is one of the largest and endangered members of family Bovidae in the Indian subcontinent [1]. Historically they were distributed across Europe to South and Southeast Asia, but currently are restricted to India, Nepal, Bhutan, Thailand, Cambodia and Myanmar [2]. The species is regionally extinct from Bangladesh, Indonesia, peninsular Malaysia, Sri Lanka, Viet Nam, Sumatra, Java and Borneo [2, 3]. With an estimated global current population size of ~4000 individuals (>2500 mature individuals) and a range-wide declining population trend, they are considered as ‘Endangered’ by IUCN [2]. The species is provided the highest protection in India and listed under the Schedule I of Wildlife Protection Act of India (1972), and in Appendix I of CITES [2]. The global population has declined significantly in recent time due to increased anthropogenic impacts and changing land use practices [2]. The long-term viability of the species is under serious threat from habitat loss and degradation, hunting, disease from livestock and hybridisation with the domestic buffalos [4]. India retains more than 90% (~3500-3700 individuals) of the global wild water buffalo population, mostly distributed as fragmented populations in two states: Assam and Chhattisgarh [5]. Majority of the population is found in northeastern state of Assam (~3000-3500 individuals) within Kaziranga National Park, Manas National Park, Dibru-Saikhowa Wildlife Sanctuary and few other areas, with a potentially small population in Meghalaya [2]. The central Indian population in the state of Chhattisgarh are clustered across a few protected areas: Udanti Wildlife Sanctuary, Bhairamgarh Sanctuary, Pamed Wildlife Sanctuary and Indravati National Park, where ~50-70 individuals are found [1]. However, the fragmented nature of the populations and the hybridisation pressure from sympatric domestic buffalos create serious challenges in their population recovery [2].

One of the main challenges to assess water buffalo population is the difficulty in morphologically distinguishing true wild buffalo from the other buffalos (free-ranging domestic buffaloes, feral buffaloes and hybrids). Throughout their distribution, the species is sympatric with the domestic buffalo and probably hybridize due to wildlife-livestock interaction [2]. As the hybrids are morphologically indistinguishable, few genetic studies have focused on identifying pure wild water buffalo populations. Preliminary genetic data (covering multiple partial mtDNA fragments) from Indian populations suggested that there is a genetic difference between wild and domestic/feral buffaloes in Kaziranga National Park, Assam and that this population is related to the population found in central India [6]. However, this data was not strong to assess the relationship between these two populations or to identify hybrids in the wild.

In this study, we sequenced the complete mitochondrial DNA of three pure wild water buffalo individuals from Assam (n=1) and Chhattisgarh (n=2), and annotated the mitogenome. We compared this mitogenome with domestic buffalo to identify sequence variations and conducted comparative analyses of secondary tRNA structures between them. Further, we confirmed the taxonomic status of wild water buffalo within the bovidae family. We feel that the results will provide baseline information to address phylogeny and identify hybrids of this endangered bovid, and will help in their conservation across their range.

## 2. Methods

### 2.1 Sampling and DNA extraction

Two tissue samples from naturally dead pure wild buffalo were received from Udanti Wildlife Sanctuary, Chhattisgarh and one from Kaziranga National Park, Assam. The samples were collected in 100% ethanol and stored in −20°C freezer till processing.

DNA was extracted from muscle tissues using QIAamp DNA Tissue Kit (QIAGEN Inc., Hilden, Germany) following the standard protocols. In brief, about 20 mg of tissue was macerated with sterile blade and digested with 30 μl of Proteinase K (20mg/ml) and 300 μl of ATL buffer (Qiagen Inc., Hilden, Germany) overnight at 56°C, followed by Qiagen DNeasy tissue DNA kit extraction protocol. DNA was eluted twice in 100 μl 1X TE buffer. We included one extraction negative during extraction to monitor possible contamination. Postextraction, one μl of eluted DNA was electrophoresed in a 0.8% agarose gel to check DNA presence and subsequently the elutions stored at −20°C freezer till further downstream processing.

### 2.2 PCR amplification and mitogenome sequencing

We amplified the whole mitogenome as overlapping fragments using bovid-specific primers described in Hassanin et al., 2009 [7]. A total of 23 different primer sets were used to amplify the mitogenome (see Supplementary Table 1 for details). PCR reactions were performed for all primers in 10μl reaction containing 4μl Qiagen multiplex PCR buffer mix (QIAGEN Inc., Hilden, Germany), 0.3 μM primer mix for each set, 4 μM BSA and 2 μl of wild buffalo DNA (1:20 dilution). PCR conditions included an initial denaturation (95 °C for 15 min); 45 cycles of denaturation (95 °C for 30 s), annealing (54-55 °C for 40 s) (See Supplementary table 1) and extension (72 °C for 40 s); followed by a final extension (72 °C for 20 min). During reactions, PCR and extraction negatives were included to monitor contamination. PCR products were visualized with 2% agarose gel. Some of the primers from Hassanin et al., 2009 [7] did not show any amplification during standardization. To cover these gaps, we designed overlapping primers by aligning *Bubalus arnee* and *Bubalus bubalis* mitogenome and using the conserved regions in these gaps (Supplementary Table 1).

Further, the amplified products were cleaned using Exonuclease-Shrimp Alkaline Phosphatase (Exo-SAP) mixture (New England Biolabs, Ipswich, Massachusetts) and sequenced bidirectionally using BigDye v3.1 terminator kit in ABI 3500XL Genetic Analyzer (Applied Biosystems, California, United States).

### 2.3 Analysis

#### 2.3.1 Mitogenome alignment and annotation

All wild buffalo sequences (n=26) were checked manually and cleaned for any nucleotide ambiguities and aligned in Mega v7 [8]. The overlapping regions were aligned to generate a complete mitogenome sequence. We used MITOS2 web [9] to annotate the consensus sequence for coding regions and their positions, incomplete stop codons and overlaps. We generated the wild buffalo mitogenome map using OGDRAW [10]. Further, we calculated nucleotide percentages and codon usages using MEGA v7 [9] and created a heat map using the ‘pheatmap’ package [11] in RStudio v1.1.383 [12]. Skew analysis was done using the following method: GC skew = (G-C)/(G+C); AT skew = (A-T)/ (A+T) to check the bias in nucleotide composition.

Finally, we predicted and visualized the secondary tRNA structures of wild buffalo using tRNAScanV [13] and compared them with the tRNA structures of domestic buffalo (*Bubalus bubalis*) to identify differences, if any. The vertebrate mitochondrial genetic code was used as a default while predicting the tRNA structures.

#### 2.3.2 Genetic differentiation with domestic buffalo and phylogenetic analyses

We estimated the mean pairwise genetic distance between the central Indian and northeast Indian wild buffalo sequence as well as with the domestic buffalo using MEGA v7 [8]. Further, we downloaded nine bovid mitogenome sequences (see Table 1) to ascertain wild buffalo phylogenetic position. A sequence of common Eland (*Tragelaphus oryx*) was used as outgroup in this analysis. We used a Bayesian approach implemented in program MRBAYES v3.2 [14]. We used a two-parameter substitution model (GTR+G, determined by jModelTest [15] and a gamma distribution of evolutionary rates across sites, with the value of the shape parameter estimated from the data. The MCMC analysis used 2 runs of 4 chains each for a run length of 1 million with sampling after every 1000 iterations till the split frequencies were below 0.01. No molecular clock was enforced in this analysis, thereby tree and branch lengths depicted topology rather than distance. We compared four independent Bayesian trees prepared with following data: a) with the complete mtDNA sequences; b) only with the protein coding genes; c) with complete Cytochrome b gene and finally d) with Cytochrome oxidase I gene. This was done to assess the best data for species-level identification, particularly between wild and domestic buffalo.

**Table 1:** Details of species mitogenome used in this study and their nucleotide composition indices.

| Scientific name | Common name | Genbank<br>accession<br>number | Whole mitogenome |  |  | Protein coding region<br>(PCG) |  |  | rRNA genes |  |  | tRNA genes |  |  | Control region |  |  |
| --- | --- | --- | --- | --- | --- | --- | --- | --- | --- | --- | --- | --- | --- | --- | --- | --- | --- |
|  |  |  | Length<br>(bp) | AT<br>Skew | GC<br>Skew | Length<br>(bp) | AT<br>Skew | GC<br>Skew | Length<br>(bp) | AT<br>Skew | GC<br>Skew | Length<br>(bp) | AT<br>Skew | GC<br>Skew | Length<br>(bp) | AT<br>Skew | GC<br>Skew |
| *Bubalus arnee*<sup>1</sup> | Wild buffalo | MT409695 | 16360 | 0.113 | -0.314 | 11361 | 0.052 | -0.325 | 2525 | 0.232 | -0.109 | 1514 | 0.034 | 0.055 | 927 | 0.054 | -0.280 |
| *Bubalus arnee*<sup>2</sup> | Wild buffalo | MT409696 | 16360 | 0.113 | -0.314 | 11361 | 0.085 | -0.376 | 2525 | 0.235 | -0.111 | 1514 | 0.034 | 0.050 | 927 | 0.055 | -0.272 |
| *Bubalus bubalis* | Domestic buffalo | AF547270.1 | 16359 | 0.113 | -0.314 | 11361 | 0.085 | -0.374 | 2525 | 0.239 | -0.118 | 1514 | 0.034 | 0.050 | 927 | 0.062 | -0.300 |
| *Syncerus caffer* | African buffalo | NC020617.1 | 16359 | 0.113 | -0.322 | 11361 | 0.087 | -0.390 | 2523 | 0.225 | -0.103 | 1514 | 0.030 | 0.062 | 931 | 0.087 | -0.390 |
| *Bos primigenius* | Auroch | GU985279.1 | 16338 | 0.102 | -0.320 | 11361 | 0.076 | -0.387 | 2526 | 0.234 | -0.102 | 1511 | 0.019 | 0.069 | 908 | 0.071 | -0.286 |
| *Bos taurus* | Cattle | NC006853.1 | 16338 | 0.102 | -0.315 | 11361 | 0.077 | -0.383 | 2525 | 0.231 | -0.096 | 1511 | 0.016 | 0.075 | 910 | 0.066 | -0.286 |
| *Bos javanicus* | Banteng | NC012706.1 | 16339 | 0.102 | -0.316 | 11361 | 0.077 | -0.383 | 2525 | 0.233 | -0.099 | 1511 | 0.016 | 0.075 | 910 | 0.065 | -0.284 |
| *Bos gaurus* | Gaur (Indian bison) | MK770201.1 | 16367 | 0.107 | -0.323 | 11361 | 0.083 | -0.392 | 2526 | 0.224 | -0.102 | 1510 | 0.030 | 0.049 | 918 | 0.063 | -0.310 |
| *Bison bison* | American bison | GU947006.1 | 16320 | 0.107 | -0.325 | 11361 | 0.081 | -0.392 | 2525 | 0.232 | -0.103 | 1513 | 0.025 | 0.066 | 890 | 0.044 | -0.293 |
| *Bos mutus* | Wild yak | KR106993.1 | 16322 | 0.105 | -0.323 | 11361 | 0.081 | -0.391 | 2527 | 0.227 | -0.103 | 1511 | 0.017 | 0.077 | 892 | 0.062 | -0.294 |
| *Bos grunniens* | Domestic yak | KR011113.1 | 16322 | 0.105 | -0.323 | 11361 | 0.081 | -0.394 | 2527 | 0.225 | -0.100 | 1511 | 0.017 | 0.077 | 892 | 0.067 | -0.299 |

<sup>1</sup>- Sample from Assam
<sup>2</sup>- Sample from Chhattisgarh. Both samples collected in this study have identical mitogenome.

## 3. Results

### 3.1 Mitogenome organisation

We generated the whole mitochondrial genome of *Bubalus arnee* (16,357 bp length). The Genbank accession numbers for the sequences are MT409695 and MT409696 for Assam and Chhattisgarh, respectively. The nucleotide composition of wild buffalo mitogenome consisted of 26.4% T, 26.6% C, 33.1% A and 13.9% G.

Composition analyses through AT and GC skew showed a positive AT value (0.112) and a negative GC value (−0.313), indicating a AT-rich mitogenome composition of wild buffalo. The annotated genes included 13 protein-coding genes (PCGs), 22 tRNA genes, two ribosomal genes and a non-coding control region. The gene order is conserved with other bovid species, and their length and positions are presented in Table 2. Out of 37 genes and control region found in the mitogenome NADH6 and eight tRNA genes (tRNA^Gln^, tRNA^Asn^, tRNA^Ala^, tRNA^Cys^, tRNA^Tyr^, tRNA^Ser^, tRNA^Glu^ and tRNA^Pro^) were encoded on the light strand (L-strand), whereas all others (n=29) are encoded on the heavy strand (H-strand).

**Table 2.**
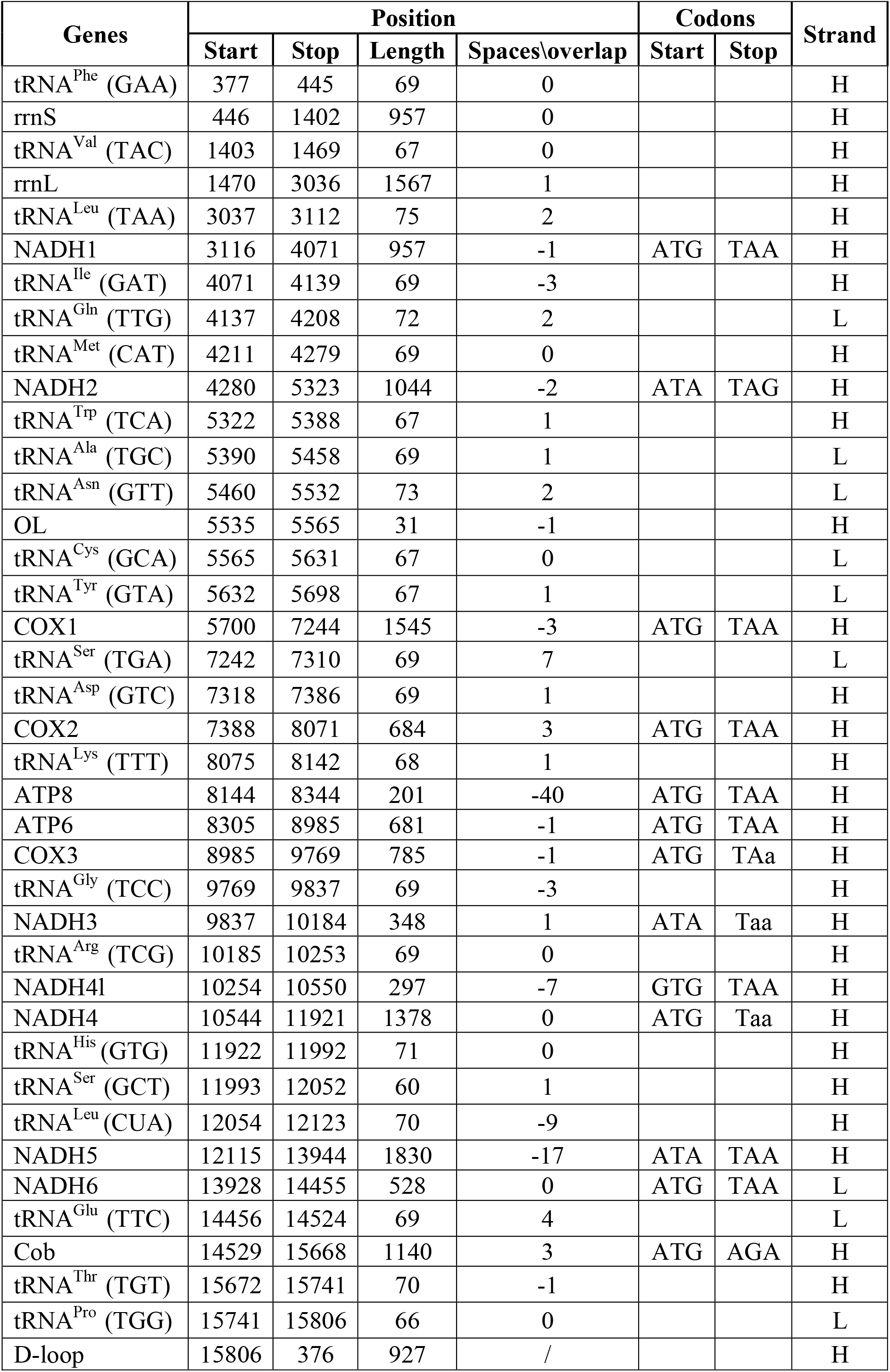
Organisation of mitochondrial genome in *Bubalus arnee*. Codons respective to each tRNA is mentioned in the first column (in parenthesis).

We observed 13 overlapping regions and 15 intergenic spaces in wild buffalo mitogenome, which are similar to other bovine species (See Table 2). Four NADH regions (NADH1, NADH2, NADH4L and NADH5), two cytochrome oxidase (COII and COIII), ATP8 and ATP6 and OL, showed overlaps ranging from 1 bp to 40 bp (ATP8 with ATP6). Three tRNA genes (tRNA^Ile^, tRNA^Thr^, tRNA^Gly^) showed the smallest overlap (1-3 bp), whereas tRNA^Lys^ gene showed the largest overlap (9 bp) with NADH5 (see Table 2). Similarly, the intergenic spaces ranged between 1-7 bp, with the longest between tRNA^Ser^ and tRNA^Asp^, respectively (see Table 2). The O_L_ was 31 bp long and found in the WANCY region between tRNA^Asn^ and tRNA^Cys^.

### 3.2 Protein Coding Genes

The total length of the wild buffalo mitochondrial protein coding genes was 11361 bp, accounting about 69.3% of the mitogenome. Average base composition was 28% T, 27.1% C, 31.1% A and 13.8% G suggesting that AT% was more than GC% in the protein coding region. The coding genes included seven NADH genes (NADH1, NADH2, NADH3, NADH4, NADH5, NADH6, and NADH4L), two ATPase (ATP6 and ATP8), three cytochrome c oxidases (COI, COII and COIII) and one cytochrome b (Cyt b). These proteincoding regions were AT skewed (Table 1). Majority of these genes (n=12) were present in the H-strand. Out of these 13 genes, NADH2, NADH3 and NADH5 have ATA as start codon whereas all others started with ATG. Similarly, most of the genes (n=9) have TAA as a stop codon except Cyt b (AGA) and NADH2 (TAG), while NADH3, NADH4 and COIII have incomplete stop codons (T at the 5’ terminal of the adjacent gene). The Relative Synonymous Codon Usage (RSCU) for the 13 wild buffalo protein-coding genes consists of 3785 codons (presented as a heatmap in Figure 3).

### 3.3 Ribosomal RNA (rRNA) and transfer RNA (tRNA) genes

The wild buffalo mitogenome has two rRNA genes: 12s rRNA and 16s rRNA, that are located between tRNA^Phe^ and tRNA^Val^, and tRNA^Val^ and tRNA^Leu^, respectively (Figure 1). Total sequence length of ribosomal RNA was 2525 bp with base composition of 37.2% A, 23.2% T, 17.6% G and 21.9% C and a positive AT skew (Table 1).

**Figure 1:**
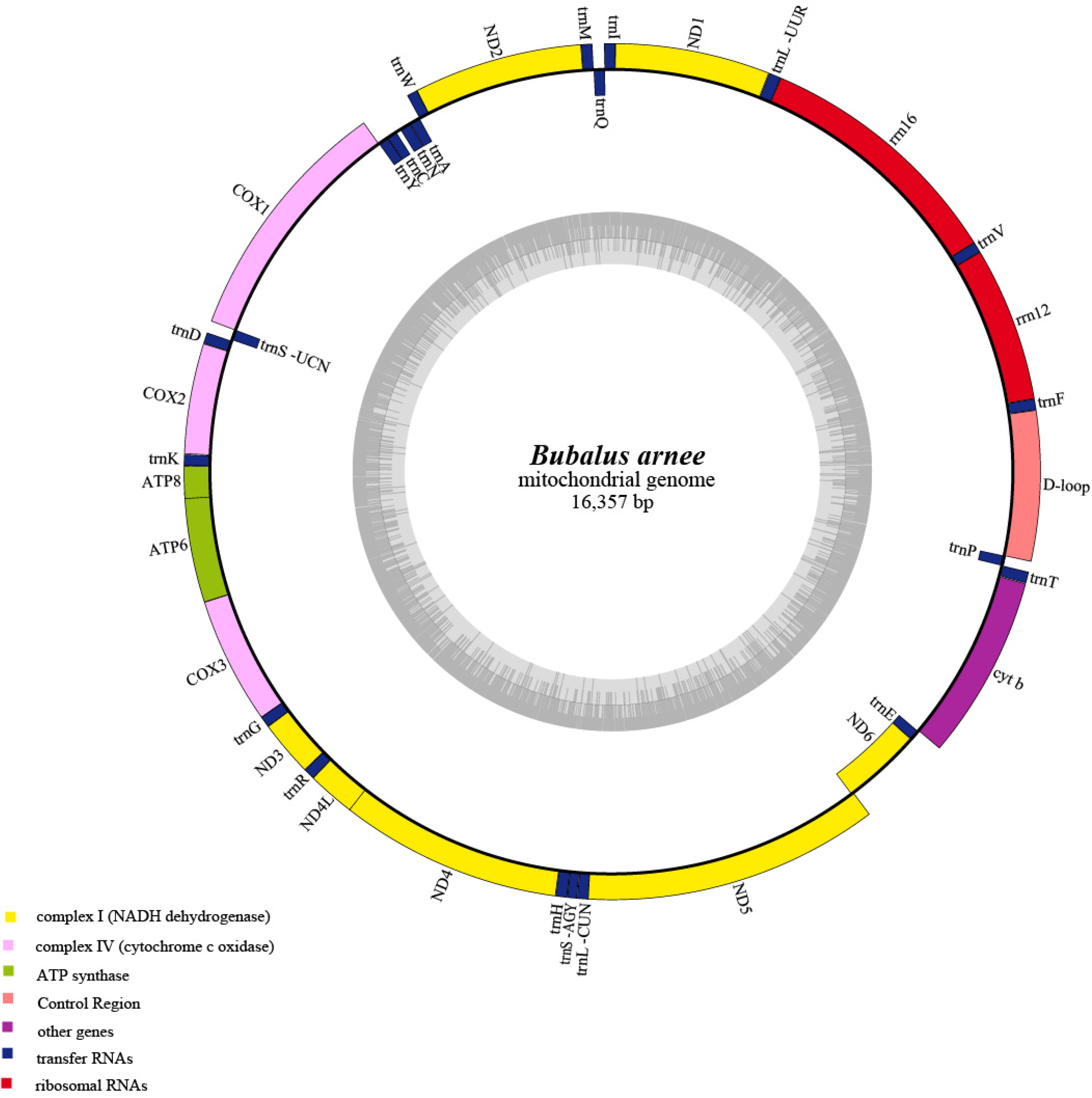
Complete mitochondrial genome of wild water buffalo (*Bubalus arnee*) with location of genes. The genes in the Heavy strands are shown outside.

**Figure 2:**
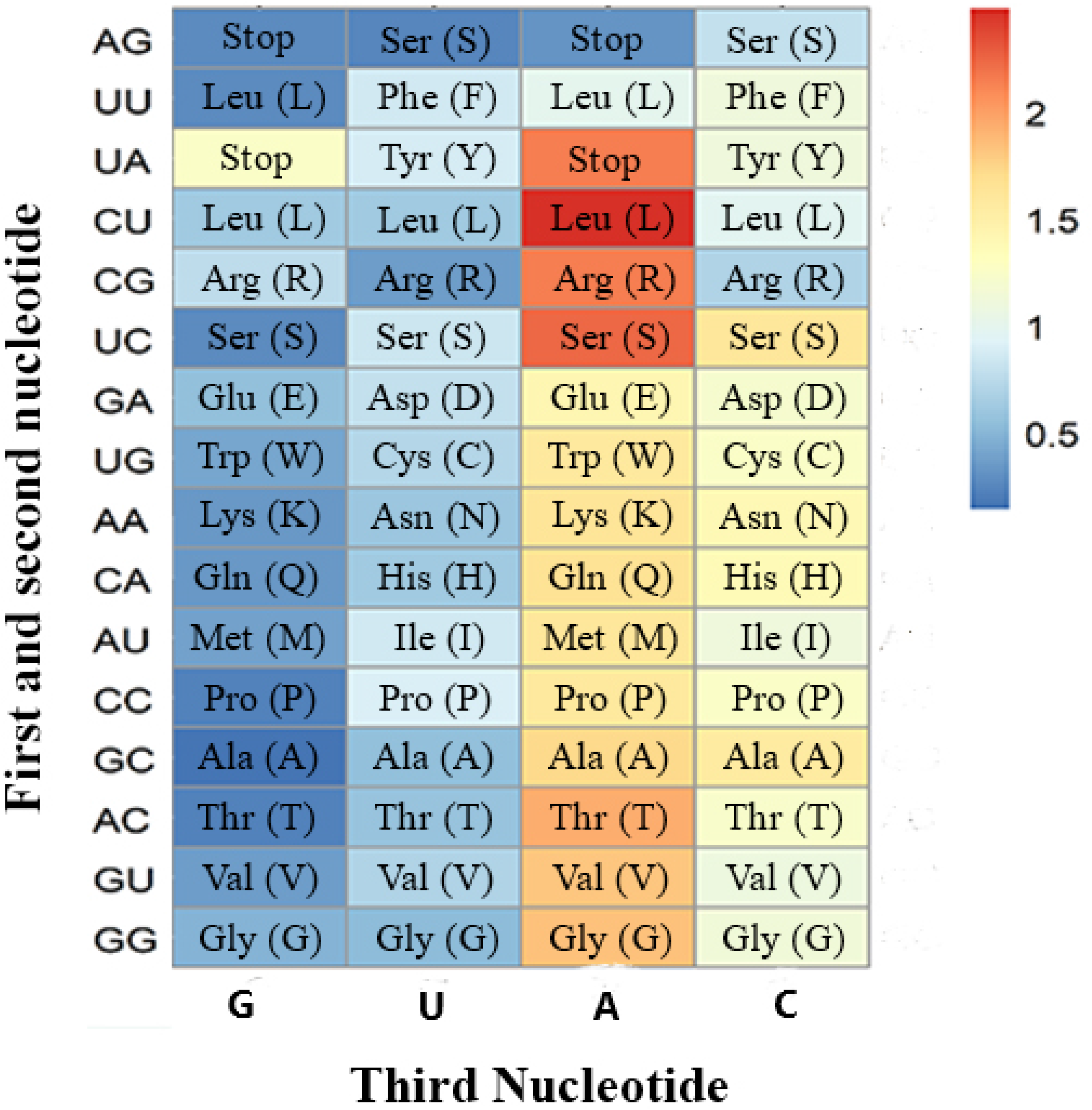
Heatmap depicting codon count to represent Relative Synonymous Codon Usage (RSCU) for the protein-coding region of *Bubalus arnee* mitochondrial genome.

The mitogenome also contains 22 tRNA genes covering a total length of 1514 bp (Table 1). Average base composition was 32% A, 29.9% T, 20.1% G and 18.1% C. Their length varied from 60bp (tRNA^Ser^) to 75 bp (tRNA^Leu^) with Leucine and Serine having two tRNAs each. Out of the 22 genes, 14 were located on H-strand and rest eight were found on L-strand. Majority of these 22 tRNA have typical secondary cloverleaf structures (n=21), while tRNA^Ser^ (GCT) lacks the D-paired arm.

Comparative structure analyses between the wild and domestic buffalo tRNA revealed that seven of the 22 tRNAs (tRNA^Trp^, tRNA^Asp^, tRNA^Thr^, tRNA^Arg^, tRNA^Lys^, tRNA^Ala^, tRNA^Leu^ (TAG)) have sequence differences between them (Figure 3) in the D-loop, TC-loop, central loop or one of the stems. Further, the tRNA^Trp^ (TCA) has sequence differences in more than one loop (Figure 3) compared to the domestic buffalo. While these differences were found in both Assam and Chhattisgarh samples, the Assam individual had one base pair difference in the tRNA^Leu^ (TAG) sequence when compared to the Chhattisgarh sample. All anticodon sequences were conserved between wild and domestic buffalo.

**Figure 3:**
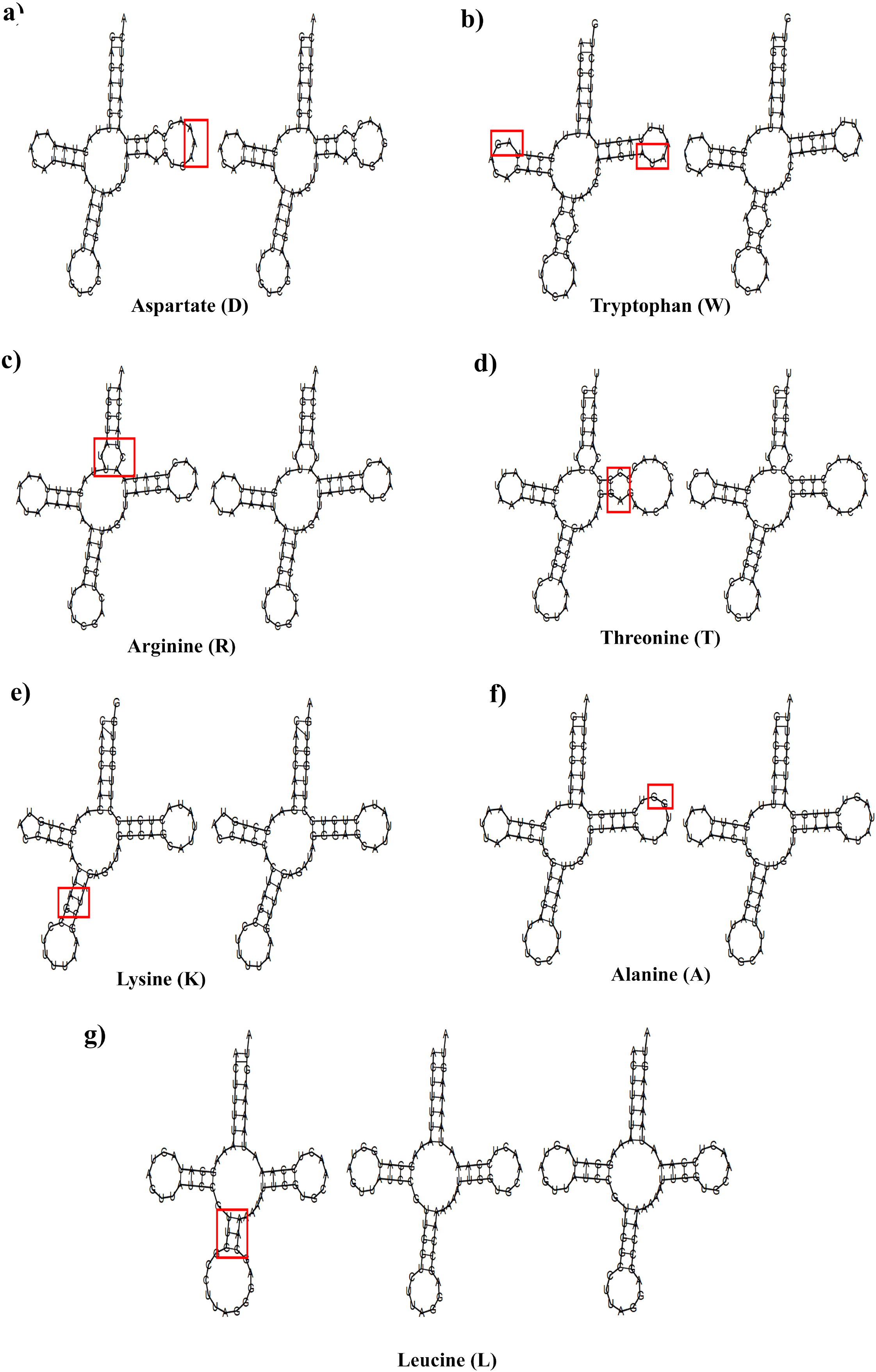
Secondary structures of seven tRNA where differences between *Bubalus arnee* and *Bubalus bubalis* (left and right pictures in all cases, respectively) have been observed. The differences are marked with red boxes.

### 3.4 Control region

The Control region (CR) of wild buffalo is 928 bp long and located between tRNA^Pro^ and tRNA^Phe^. The base composition of CR was 28.3% T, 25.8% C, 31.5% A and 14.5% G. The average AT and GC skew was 0.053 and −0.28, respectively (Table 1). The control region length varied considerably across the bovid species used in this study, ranging between 890 bp in bison (*Bison bison*) to 931 bp in Cape Buffalo (*Syncerus caffer*) (Table 1). Such INDELs have been found across many mammalian species [16, 17].

### 3.5 Genetic distance between wild and domestic buffalo

Both the Chhattisgarh wild buffalo mitogenome were identical. The mitogenomes of wild (Assam and Chhattisgarh samples) and domestic buffalo showed 384 variable sites across the mitogenome. Majority of these sites (n=275, 71.61%) were found within the coding region, including 211 synonymous mutations, 64 replacement changes and no INDELs. The nucleotide diversity between wild and domestic buffalo was 0.01565 and the average number of nucleotide difference (k) was 256. However, between the Assam and Chhattisgarh samples we found only 17 polymorphic sites and nucleotide diversity was 0.00104 (15 polymorphic sites and nucleotide diversity of 0.00097 in the coding region), indicating that they are genetically much closer than the domestic buffalo. Genetic distance between the wild buffalo samples from Assam and Chhattisgarh was 0.001 whereas between domestic and wild buffalo was 0.024 (0.022 with coding regions), suggesting much higher genetic differences between wild and domestic buffalos.

### 3.6 Phylogenetic analysis

Earlier phylogenetic research conducted suggested that the riverine and swamp buffalo was domesticated from wild buffalo in the Indian peninsula [18]. Flamand et al. (2003) in their microsatellite-based work has focused on detecting hybrids and found that earlier studies might have used hybrid individuals as pure for analyses [4]. We constructed Bayesian trees with the whole mitogenome, coding region, COI and Cyt b to assess wild buffalo taxonomy with nine bovidae species (see Table 1). All four phylogenetic trees showed very similar phylogenetic positions for wild buffalo, domestic buffalo and African buffalo (Figure 4). They all formed a paraphyletic clade with the genus *Bos*. Within the *Bubalus* clade both Chhattisgarh and Assam sequences form a monophyletic clade, sister to *Bubalus bubalis* with high posterior probabilities (1.00) (Figure 4). The clustering pattern of Bovidae was broadly consistent with previous studies [19].

**Figure 4:**
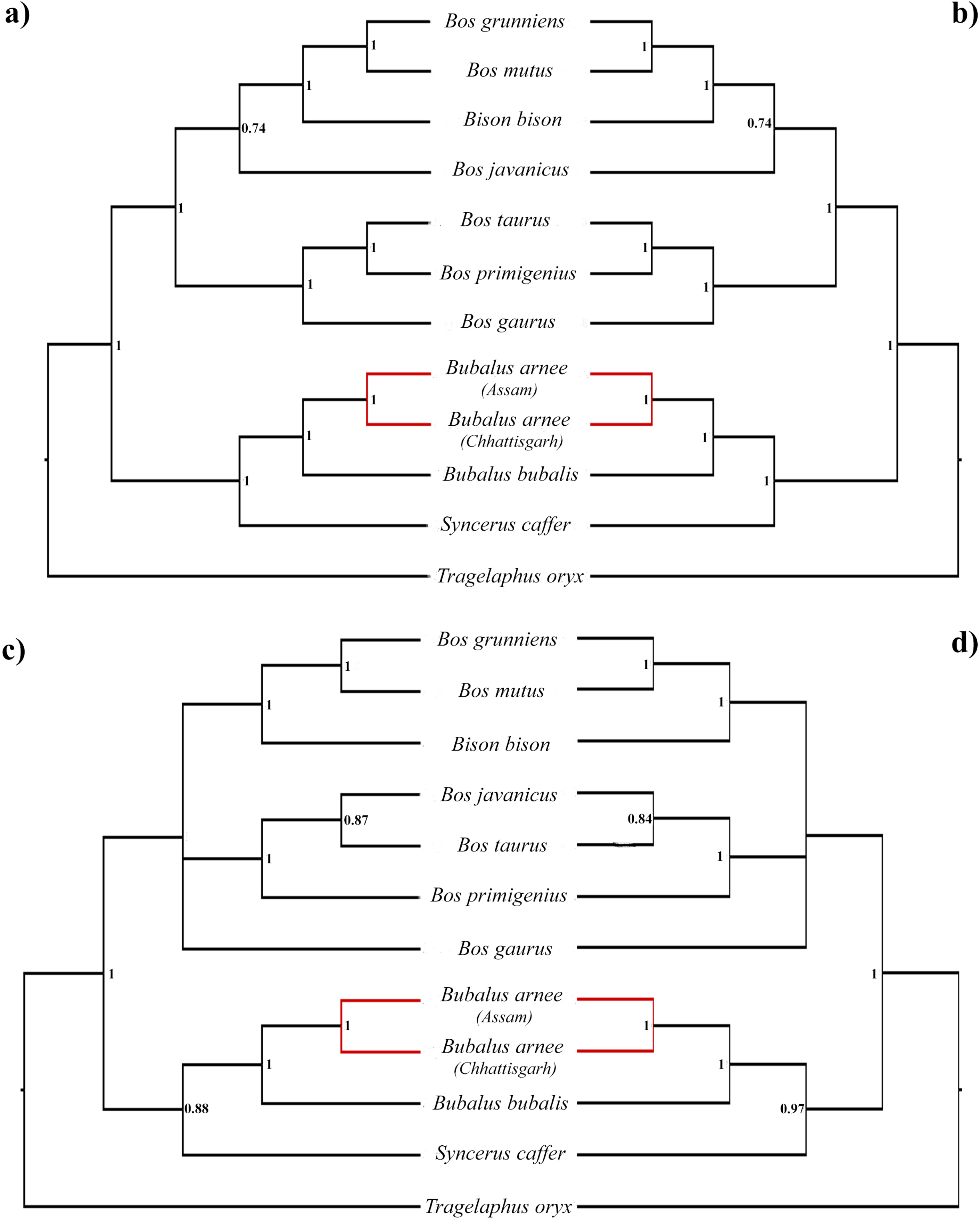
Phylogenetic relationships among wild buffalo and other bovid species with *Tragelaphus oryx* as outgroup inferred from a) whole mitogenome b)13 PCGs, 22 tRNA and two rRNA c) Cytochrome oxidase-I d) Cytochrome b using Bayesian inference (BI). Bayesian posterior probability values are shown at each node.

## 4. Discussion

To the best of our knowledge, this is the first report of complete mitogenome sequence (16,357 bp in length) and annotations of wild water buffalos from two extant populations (Assam and Chhattisgarh) in India. Both populations face serious threats from habitat loss, hunting, resource depletion, diseases and hybridization with domestic water buffalo. Latest IUCN assessments of wild water buffalo suggest that hybridization with domestic buffalo is the most critical concern that is projected to continue [2]. However, till date no comprehensive evaluation of the extent of hybridization and its effects has been conducted, possibly due to difficulties in identification of pure wild water buffalo. We collected pure wild buffalo samples from both habitats and phylogenetic analyses with complete mitogenome sequence (as well as with specific genes) showed sister clade with domestic buffalo. It is known that phenotypic and behavioural criteria are not sufficient to differentiate between wild and hybrid wild buffalo [20, 21]. Our analyses with complete cytochrome b/ cytochrome oxidase I suggest that it can separate pure wild buffalo and domestic buffalo sequences. Further, our results clearly demonstrate that secondary structures of seven tRNA are different between the wild and domestic buffalo. However, it is important to remember that only mtDNA-based assessment can be misleading to confirm species purity as it is maternally inherited [4]. A combination of mtDNA and nuclear DNA (microsatellite, SNPs etc.) based analyses will provide much stronger inferences regarding hybridization (for example, [16] for Bison and [22] for African elephants). Future studies should focus on developing/standardizing such markers for in-depth understanding of hybridization effects, with emphasis on functional implications of the tRNA structural differences.

Although our data suggest that the Assam and Chhattisgarh samples are two different haplotypes (with 17 nucleotide differences), the genetic distance is very low supporting the earlier report [7]. Given the central Indian wild buffalo population in Chhattisgarh is very small [1], the Assam population can act as potential source for reintroduction program. However, we would like to point out that a more extensive population level study combining larger, informative fragments of mtDNA and nuclear DNA is required before selecting the founders for any such reintroduction program. If the Assam population is genetically structured then appropriate selection criteria need to be developed. Finally, the complete mitochondrial DNA sequence data would also help us to develop species-specific assays for identification of wild and domestic buffalo, and help in forensic identification of seized buffalo contrabands (meat, horn, other body parts).

In conclusion, we believe that the first complete mitogenome of wild water buffalo (*Bubalus arnee*) would act as a baseline information to further extend more research on phylogeny, phylogeography and hybridization and help in conservation of this endangered species across its range. This work also emphasizes the importance of similar work on less-known species that require immediate conservation attention.

## Acknowledgement

We thank Chhattisgarh and Assam Forest Departments for providing pure wild water buffalo samples for this work. We also thank the Director, Dean and Research Coordinator of Wildlife Institute of India for their support. This study was funded by the INSPIRE Faculty Award from Department of Science and Technology, Government of India (IFA12-LSBM-47). The funders had no role in study design, data collection and analysis, decision to publish, or preparation of the manuscript.

---

**Supplementary Table 1:**
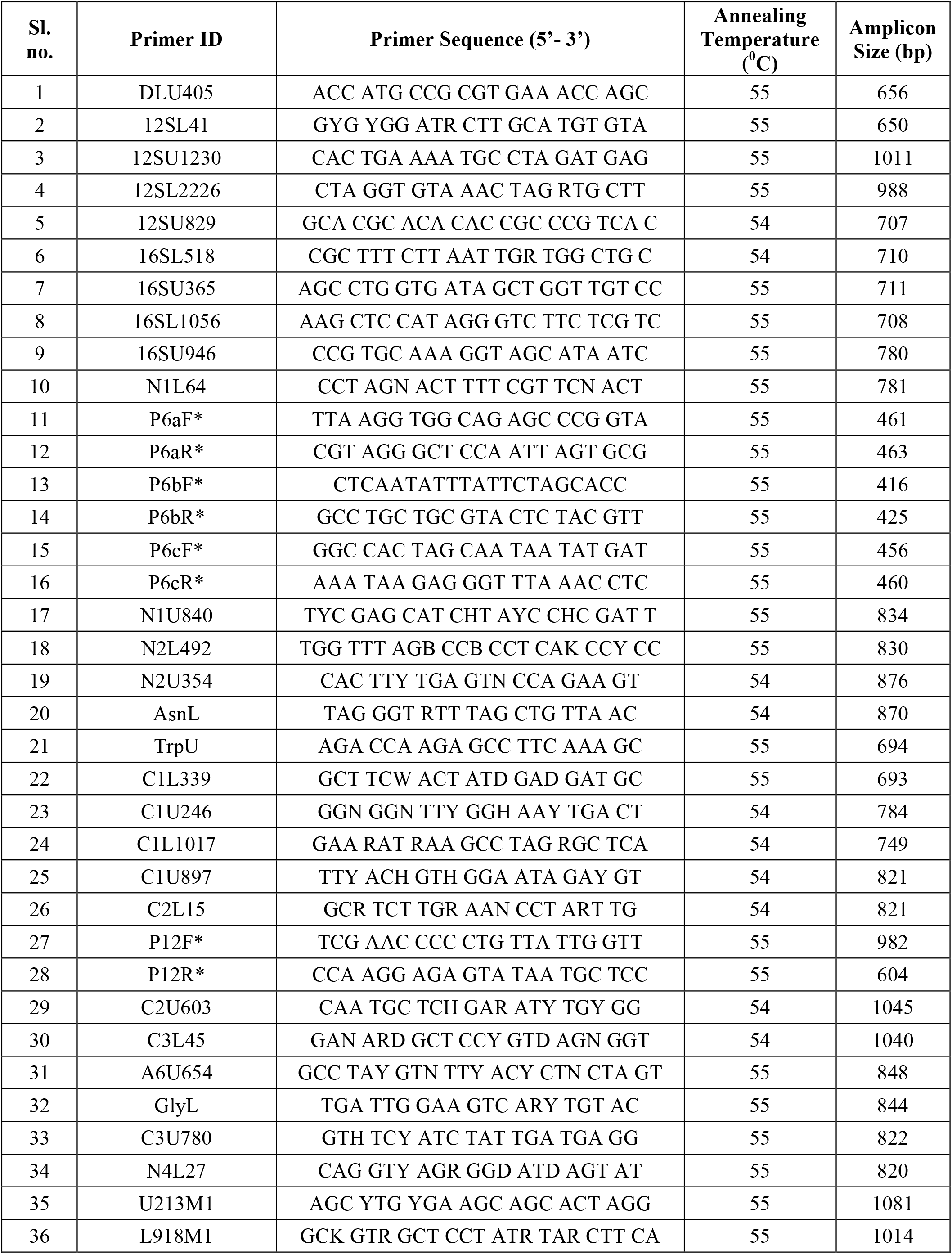
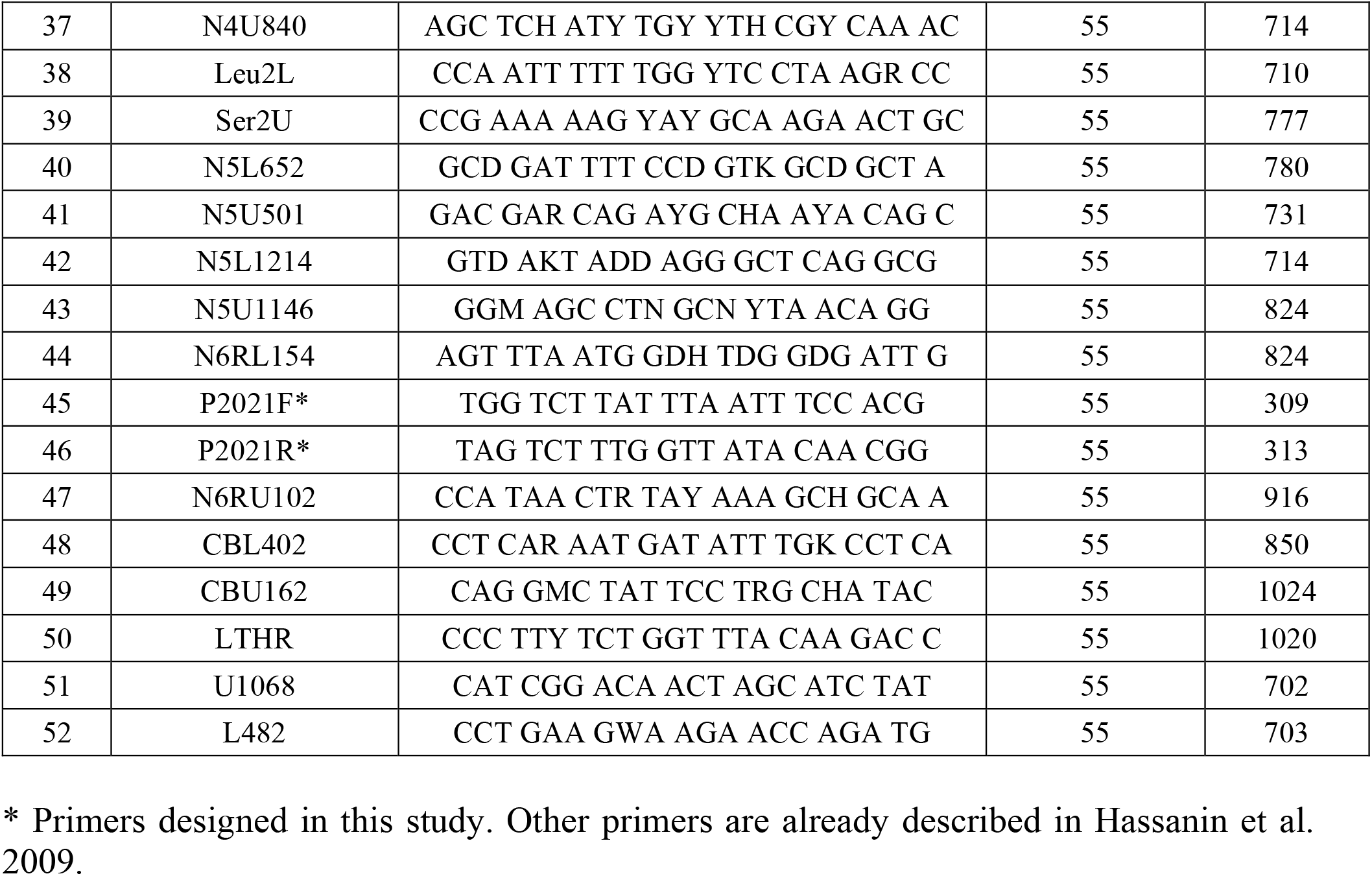
Details of the primers used in this study to amplify wild water buffalo (*Bubalus arnee*) mitogenome.

## References

[1] Ranjitsinh, M. K., Verma, S. C., Akhtar, S. A., Patil, V., Sivakumar, K., & Bhanubhakude, S. (2004). Status and conservation of the wild buffalo *Bubalus bubalis* in Peninsular India. Bombay Natural History Society, 101(1), 64–70.

[2] Kaul, R., Williams, A. C., Rithe, K., Steinmetz, R., & Mishra, R. (2019). The IUCN Red List of Threatened Species. http://dx.doi.org/10.2305/IUCN.UK.2019-1.RLTS.T3129A46364616.en

[3] Corbet, G. B., & Hill, J. E. (1992). The mammals of the Indo-Malayan region: a systematic review (Vol. 488). Oxford: Oxford University Press.

[4] Flamand, J. R. B., Vankan, D., Gairhe, K. P., Duong, H., & Barker, J. S. F. (2003). Genetic identification of wild Asian water buffalo in Nepal. Animal Conservation, 6(3), 265–270. https://doi.org/10.1017/S1367943003003329

[5] Choudhury, A. (1994). The decline of the wild water buffalo in northeast India. Oryx, 28(1), 70–73. https://doi.org/10.1017/S0030605300028325

[6] CCMB. 2010. A Brief Report on Genetics of Wild Buffaloes at Udanti Wildlife Sanctuary. Central for Cellular and Molecular Biology, Hyderabad, India.

[7] Hassanin, A., Ropiquet, A., Couloux, A., & Cruaud, C. (2009). Evolution of the Mitochondrial Genome in Mammals Living at High Altitude: New Insights from a Study of the Tribe Caprini (Bovidae, Antilopinae). Journal of Molecular Evolution, 68(4), 293–310. https://doi.org/10.1007/s00239-009-9208-7

[8] Kumar, S., Stecher, G., & Tamura, K. (2016). MEGA7: Molecular Evolutionary Genetics Analysis Version 7.0 for Bigger Datasets. Molecular Biology and Evolution, 33(7), 1870–1874. https://doi.org/10.1093/molbev/msw054

[9] Bernt, M., Donath, A., Jühling, F., Externbrink, F., Florentz, C., Fritzsch, G., Pütz, J., Middendorf, M., & Stadler, P. F. (2013). MITOS: Improved de novo metazoan mitochondrial genome annotation. Molecular Phylogenetics and Evolution, 69(2), 313–319. https://doi.org/10.1016/j.ympev.2012.08.023

[10] Greiner, S., Lehwark, P., & Bock, R. (2019). Organellar Genome DRAW (OGDRAW) version 1.3.1: Expanded toolkit for the graphical visualization of organellar genomes. Nucleic Acids Research, 47(W1), W59–W64. https://doi.org/10.1093/nar/gkz238

[11] Kolde, R. (2013) pheatmap: Pretty Heatmaps. version 0.7.4 edn. R package

[12] RStudio. 2015. RStudio: integrated development environment for R. Boston: RStudio Inc.

[13] Chan, P. P., & Lowe, T. M. (2019). tRNAscan-SE: Searching for tRNA Genes in Genomic Sequences. In M. Kollmar (Ed.), Gene Prediction (Vol. 1962, pp. 1–14). Springer New York. https://doi.org/10.1007/978-1-4939-9173-0_1

[14] Ronquist, F., Teslenko, M., van der Mark, P., Ayres, D. L., Darling, A., Höhna, S., Larget, B., Liu, L., Suchard, M. A., & Huelsenbeck, J. P. (2012). MrBayes 3.2: Efficient Bayesian Phylogenetic Inference and Model Choice Across a Large Model Space. Systematic Biology, 61(3), 539–542. https://doi.org/10.1093/sysbio/sys029

[15] Darriba, D., Taboada, G. L., Doallo, R., & Posada, D. (2012). jModelTest 2: More models, new heuristics and parallel computing. Nature Methods, 9(8), 772–772. https://doi.org/10.1038/nmeth.2109

[16] Douglas, K. C., Halbert, N. D., Kolenda, C., Childers, C., Hunter, D. L., & Derr, J. N. (2011). Complete mitochondrial DNA sequence analysis of *Bison bison* and bison–cattle hybrids: Function and phylogeny. Mitochondrion, 11(1), 166–175. https://doi.org/10.1016/j.mito.2010.09.005

[17] Kumar, A., Gautam, K. B., Singh, B., Yadav, P., Gopi, G. V., & Gupta, S. K. (2019). Sequencing and characterization of the complete mitochondrial genome of Mishmi takin (*Budorcas taxicolor taxicolor*) and comparison with the other Caprinae species. International Journal of Biological Macromolecules, 137, 87–94. https://doi.org/10.1016/j.ijbiomac.2019.06.201

[18] Nagarajan, M., Nimisha, K., & Kumar, S. (2015). Mitochondrial DNA Variability of Domestic River Buffalo (*Bubalus bubalis*) Populations: Genetic Evidence for Domestication of River Buffalo in Indian Subcontinent. Genome Biology and Evolution, 7(5), 1252–1259. https://doi.org/10.1093/gbe/evv067

[19] MacEachern, S., McEwan, J., & Goddard, M. (2009). Phylogenetic reconstruction and the identification of ancient polymorphism in the Bovini tribe (Bovidae, Bovinae). BMC Genomics, 10(1), 177. https://doi.org/10.1186/1471-2164-10-177

[20] Heinen, J.T. (2002) Phenotypic and behavioural characteristics used to identify wild buffalo from feral backcrosses in Nepal. Journal of the Bombay Natural History Society, 99, 173–183

[21] Dahmer, T. D. 1978. Status and ecology of wild Asian buffalo *(Bubalus bubalis* L.) in Nepal. M.S. Thesis, University of Montana.

[22] Mondol, S., Moltke, I., Hart, J., Keigwin, M., Brown, L., Stephens, M., & Wasser, S. K. (2015). New evidence for hybrid zones of forest and savannah elephants in Central and West Africa. Molecular Ecology, 24(24), 6134–6147. https://doi.org/10.1111/mec.13472

